# Designing meaningful continuous representations of T cell receptor sequences with deep generative models

**DOI:** 10.1101/2023.06.17.545423

**Authors:** Allen Y. Leary, Darius Scott, Namita T. Gupta, Janelle C. Waite, Dimitris Skokos, Gurinder S. Atwal, Peter G. Hawkins

## Abstract

T Cell Receptor (TCR) antigen binding underlies a key mechanism of the adaptive immune response yet the vast diversity of TCRs and the complexity of protein interactions limits our ability to build useful low dimensional representations of TCRs. To address the current limitations in TCR analysis we develop a capacity-controlled disentangling variational autoencoder trained using a dataset of approximately 100 million TCR sequences, that we name TCR-VALID. We design TCR-VALID such that the model representations are low-dimensional, continuous, disentangled, and sufficiently informative to provide high-quality TCR sequence *de novo* generation. We thoroughly quantify these properties of the representations, providing a framework for future protein representation learning in low dimensions. The continuity of TCR-VALID representations allows fast and accurate TCR clustering, benchmarked against other state-of-the-art TCR clustering tools and pre-trained language models.

## INTRODUCTION

T Cell Receptors (TCRs) are protein complexes that are present on the surface of T cells and are selected in the thymus to bind non-self peptide antigens[1]. This forms an important arm of our adaptive immune response with its ability to kill cells that have been infected or have other mutations that may cause harm, for example in cancer. It is becoming increasingly common to perform high-throughput measurements of TCRs in biological samples [2], but such data do not often include information about the antigens those TCRs bind. Though concurrent high-throughput measurements of TCRs and cognate antigens are possible [3, 4] they require prior selection of antigens for study, leading to bias. However, these technologies do provide datasets with which to train accurate classification models of TCR-antigen interactions for those studied antigens [3, 5].

There are estimated to be 10^15^ - 10^61^ possible human TCR proteins [6, 7], with approximately 10^11^ T cells in an individual, there is consequently very limited overlap of TCR sequences between individuals. There is estimated to be an even larger space of possible antigens that these TCRs can bind, such that TCR cross-reactivity is thought to be essential [8]. This incredibly large state space with non-injective interactions make it challenging to build general predictive models for TCR-antigen interactions in an antigen agnostic manner. Experimentally defined groups of TCRs with differing functional properties can lead to differing phenotypes, such as diverging cell states [9]. Tumor growth has been shown to be controlled in a way that correlates with the TCR function [10].

Clustering TCRs that are sufficiently similar to bind the same antigen has been studied in the case where the antigen is known [3]. In the antigen agnostic case, TCR clustering has been approached via sequence-based evolutionary distance metrics between TCRs [11, 12], and more recently via autoencoder [5, 13] and masked language models [11, 12].

Here we developed an “atlas” of TCRs via representation learning, to provide the ability to cluster TCRs based on these representations, classify TCRs by known ability to bind certain antigens, and to do so in an interpretable way. Further, with the advent of TCR based therapeutics [14] and the similarity of TCRs to antibodies, we designed the atlas to be generative such that the atlas provides a future route to *de novo* TCR/antibody design. Functional data for antibodies and TCRs are often limited and costly to perform, which makes Bayesian Optimization (BO) a useful tool in this space to select sequences for further design iterations [15]. Such optimization requires a low dimensional space with a continuous function being optimized, and for that reason BO in the latent space of deep autoencoders (DAEs) has gained recent interest for complex data types [16–19].

Masked language models of proteins and TCRs typically use large dimensional representations per amino acid [11, 20] which can be problematic for representations of full protein sequences [21], and are not inherently generational in contrast with DAEs and auto-regressive language models such as GPT-3 [22]. Although DAE representations of TCRs have been developed [5, 13], the continuity of their latent spaces has not been throughly investigated. Indeed, investigation of the continuity of latent spaces in general is often only studied on toy models with known generative processes [23].

The field of disentangled representation learning (DRL) aims to learn representations of high dimensional data that improve the interpretability of models and their generational capabilities. DRL approaches have been applied to TCRs [24] but none to our knowledge have fully explored the tradeoffs between the landscape smoothness, interpretability, and sequence generation.

Here we train TCR-VALID (T Cell Receptor - Variational Autoencoder Landscape for Interpretable Disentangling), a capacity controlled 𝒞*β*-Variational Auto Encoder (VAE) [25, 26] model trained using approximately 100 million unique TCRs from a combination of *α* and *β* chains and assess the ability to generate high-quality sequences from this generative model. The disentanglement of the latent space of TCR-VALID is quantitatively evaluated, and we modify continuity metrics from machine learning literature to our biological context to measure the continuity of these representations in a systematic way. We show that the underlying assumption that TCRs with similar sequences will be embedded closely in our space provides state of the art TCR clustering, which together with continuity of the space provides a future route to BO in this low dimensional space.

## RESULTS

### Unsupervised learning of a TCR landscape via physicochemical feature embedding

Though the state space of TCR-antigen interactions is very large and we have very few measured interaction pairs involving a small subset of antigens, we do have larger volumes of TCR sequence data in the absence of known antigen pairings. We therefore hypothesized that using the unlabeled TCR sequence data to build latent representations of TCRs could then be used for downstream clustering, classification, and *de novo* generation. TCRs are made of two subunits, which for the majority of TCRs are *α* and *β* TCR chains. Although these chains occur in distinct pairs, single cell sequencing is required to resolve the pairing and thus the datasets of paired TCR chains are much smaller than datasets of independent *α* and *β* chains. Consequently, we chose to model TRA and TRB sequences independently in order to get a larger sampling of the state space of each sequence type during training.

The interaction of TCRs and antigen is primarily encoded by the Complimentary Determining Regions (CDRs) [1, 27] of the TCR chains. The first two CDRs, CDR1 and CDR2, are encoded uniquely by a given V gene, whereas the CDR3 region occurs at the site of V(D)J recombination [1] and includes quasi-random nucleotide insertions and deletions. This leads to TCR chains being the product of a sparse discrete space for the V gene selection and a denser discrete space for the CDR3. This type of problem occurs often in biological sequence data where there is a sparse family structure that is complicated by dense variation at the sub-family level.

TCRs that bind the same antigen show biases to certain V gene usage [3], and so it is essential that information about V gene is encoded into the latent space for TCR clustering and classification. We sought to build a latent space that encodes the sparse discrete V genes and the highly variable CDR3 sequences in the same space such that we can co-cluster TCRs with similar yet distinct V genes, thereby precluding conditional-VAE models. The V gene can almost be uniquely encoded by the CDR2 sequence, so we chose to use the amino acid sequence of the CDR2 joined by a gap character to the the CDR3 to describe a single TCR chain (Fig.1a). This is beneficial for encoding the biophysical similarity of two V genes, rather than relying on a dictionary embedding strategy [5]. The structural interaction of the CDR loops with the antigen is dictated by local physical interactions driven by the physicochemical properties of the amino acids of the loops. To capture this, we encoded the amino acids into 7 physicochemical features (see methods) forming a 2-D description for each TCR, and sought to learn smooth, low dimensional representations of these physicochemically encoded TCRs.

**FIG. 1.**
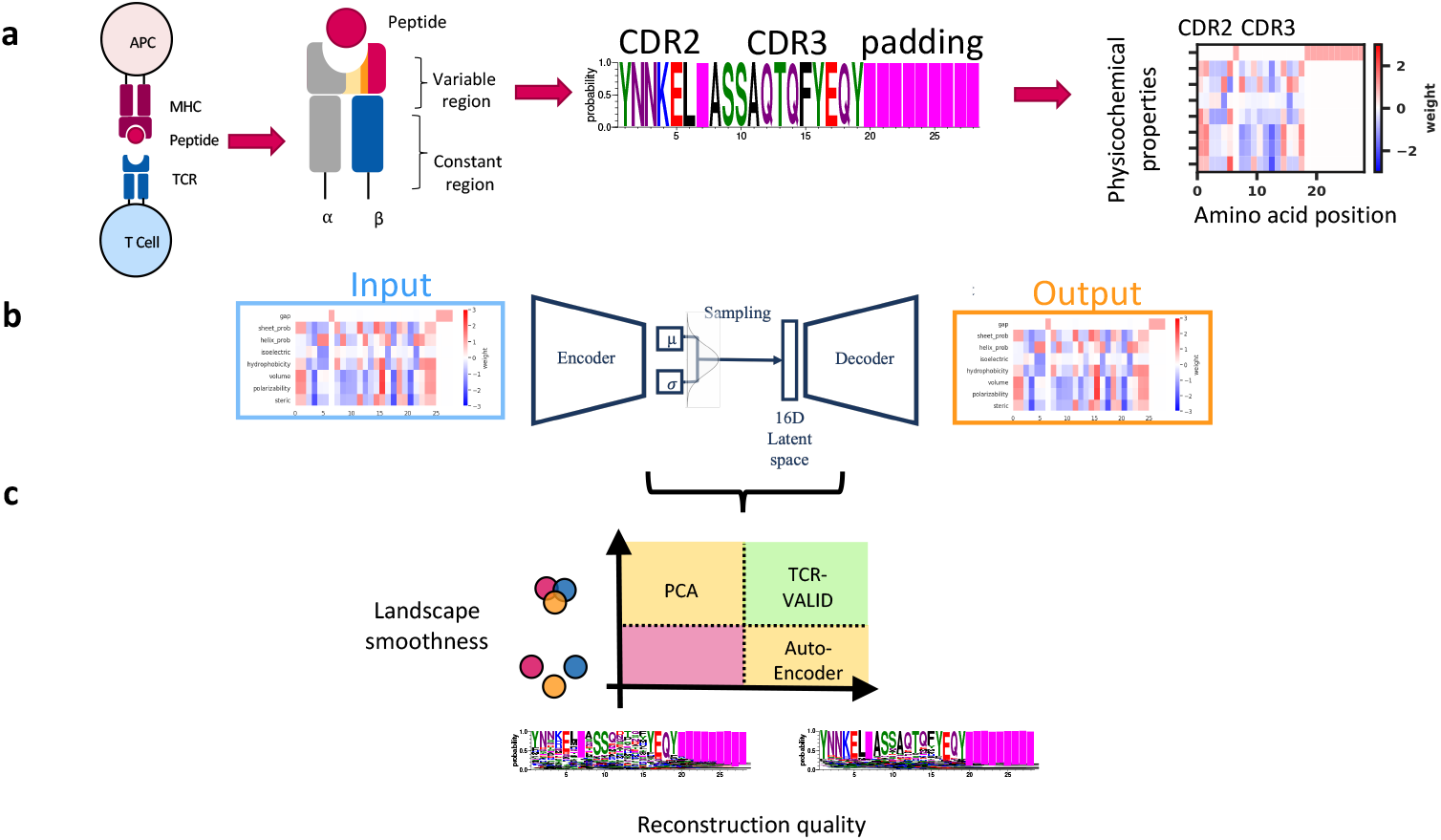
TCR-VALID: Learning a physicochemically informed latent landscape for TCR sequences: TCRs interact with peptide:MHC antigens on Antigen Presenting Cells (APC) primarily via the CDR loops of their variable region (a), since the V gene usage can be encoded almost uniquely via CDR2 and remaining diversity is in the CDR3, we use these sequence regions to encode a single TCR chain, and subsequently use physicochemical features of the amino acids to represent the sequence. (b) TCR-VALID architecture diagram: physicochemically encoded TCR sequences are used as input to train a capacity controlled 𝒞*β* VAE to learn a continuous 16D latent representation of TCR sequences. (c) Diagram illustrating the tradeoffs between representation learning approaches for TCR sequences. TCR-VALID aims to balance those tradeoffs in order to provide learned landscape smoothness, interpretability without compromising TCR sequence reconstruction quality.

We chose to base our TCR-VALID architecture (Fig.1a) on a capacity-controlled disentangling 𝒞*β*-VAE [26], using a dataset of approximately 100 million TCR sequences (methods: data). Given our reasonably short sequences, we were able to make use of light-weight convolutional neural networks (CNNs) for the encoder and decoder to learn the highly non linear patterns that underlie the the key features of a given TCR. We explored the properties of the latent representations of TCRs (Fig.1c) and the ability of the decoder to generate physicochemical representations of TCRs in a continuous space which ultimately form position weight matrices (PWMs) of TCRs (Fig.1b, methods).

### Balancing landscape continuity with sequence reconstruction accuracy

Our VAE models on the physicochemical representations of TCRs generate a continuous space of PWMs. This allows for small motions in latent space to slowly change the properties of the generated TCR sequence rather than discretely change specific amino acids. However, the sparse structure of the TCR space driven by the relatively small number of V genes can generate large regions of discontinuity in the latent space. This can cause challenges when clustering TCRs which may be similar in their CDR3 and V gene physicochemical properties, yet are distinct entities. It can also cause challenges in generation of TCRs due to regions of the latent space that do not correspond to the true manifold of physicochemically feasible TCRs.

For *β*-VAEs, one tunes the relative weight of the reconstruction loss and Kullback-Leibler divergence that forces the multivariate normal distribution on the latent space, where *β* = 1 corresponds to a standard VAE. Importantly, due to the TCR physicochemical representations being continuous, one cannot measure the exact bits of difference between the input and reconstruction, meaning that *β* = 1 does not carry the same meaning as in a typical VAE. The capacity controlled VAE [26] allows one to control the quantity of information in the latent space via the capacity term, 𝒞. This allows us to tune the smoothness of the latent space in a principled way, larger 𝒞 leading to better reconstruction at the expense of landscape continuity (Fig.2a). We found this capacity limiting approach aids in preventing posterior collapse in the low 𝒞 regime. The capacity term is simpler to implement and interpret than other methods of controlling the information in the latent space that rely on dynamically tuning the terms of the training loss [29, 30].

**FIG. 2.**
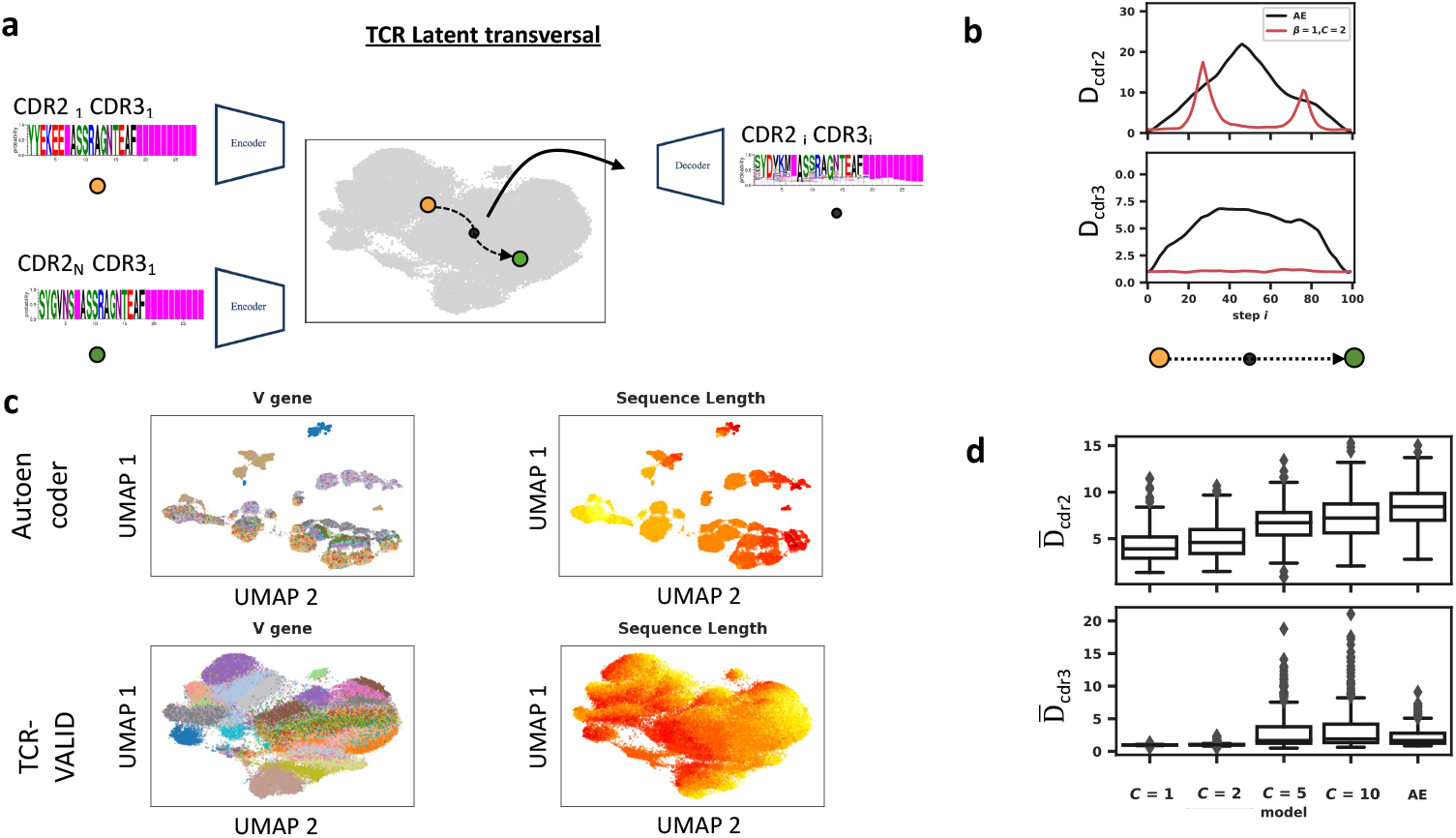
TCR-VALID learns a smooth and realistic embedding of TCRs. (a) Illustration of TCR latent transversals to evaluate embedding landscape smoothness: two TCR sequences (CDR2-3) with identical CDR3 but differing CDR2 are embedded into the learned latent space. A linear interpolation between TCRs in latent space is then decoded back into sequence space and evaluated. (b) The distance between TCR-VALID (red) and auto-encoder (black) decoded interpolated TCRs and training TCR sequences for CDR2 (*D*_*cdr*2_ metric, top) and CDR3 (*D*_*cdr*3_, bottom), (c) UMAP embedding for subset of TCR training sequences for auto-encoder (top) and TCR-VALID (bottom) colored by V gene usage (left panel) and sequence length (right panel). (d) Averaged distance over the whole trajectory for many Monte Carlo selected latent space transversals for both CDR2 (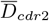 metric, top) and CDR3 (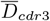 metric, top).

We sought to quantitatively evaluate the continuity of latent TCR landscapes as a function of the information stored in the latent space. We aimed for all points between any pair of TCRs in the latent landscape to smoothly represent the manifold of feasible TCRs, driven by both discrete V genes and dense CDR3 sequences. Inspired by the latent transversals employed to evaluate simple image datasets [23, 26], we developed a TCR latent transversal metric (methods). Briefly, we randomly select two unique TCR sequences with identical CDR3s and different CDR2s and embed them into our latent landscape with our trained encoder, and linearly interpolate between those two embedded TCRs in the latent space generating TCR PWMs along the trajectory (Fig.2a). The distance *D*_*cdr*2_ measures the difference between the CDR2 of the generated TCR and the closest observed CDR2 in the reference library, thereby measuring proximity to the true data manifold such that increases in this distance along a traversal indicate discontinuities in the latent landscape. Additionally we measure *D*_*cdr*3_, which is the distance from the interpolated decoded CDR3 along the traversal to the CDR3 of the two endpoint TCRs. In order to not penalize models with lower ability to encode information due to lower capacity term 𝒞, which will systematically have worse reconstruction accuracy even at the traversal endpoints, we normalize *D*_*cdr*3_ to the distances endpoints (details in methods). We do not expect CDR3 to change along the trajectory and thus expect *D*_*cdr*3_ to be small for a smoothly encoded latent space.

In order to balance the landscape smoothness with the amount of information the 𝒞*β*-VAE latent landscape can encode we tuned the capacity 𝒞 on a randomly sampled reduced dataset of ∼4 million TRB chains. For the lowest 𝒞 of 1 nat per latent dimension, we required a greater weighting of the capacity control term *β* in the loss to reach the capacity (methods). Analyzing the average distances 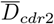 and 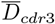 over many Monte Carlo selected latent space traversals for a range of latent space information capacities we find that capacity of 2 or fewer nats per dimension leads to smooth, continuous latent representations (Fig.2e). By visualizing the latent trajectories for an autoencoder and a hyperparameter-optimized TCR-VALID model with capacity of 2 nats per dimension, we can see that the learned latent landscape strays considerably further from the manifold of physicochemically feasible TCRs for autoencoders (Fig.2d) as is reflected in the averaged 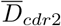 and 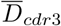 metrics.

### Latent dimensions are disentangled and allow for principled TCR generation

Another key objective of TCR-VALID is to provide an interpretable TCR landscape which would enable characterization of the manifold of TCR sequences. Interpretability of TCR latent space can be defined as the ability to encode biological and physicochemical intuition and reasonability into our model output [31]. Unlike many datasets used to quantitatively benchmark disentangling properties, not all the ground truth generative factors for TCR sequences are known, labeled, or independent.

TCRs are generated biologically via the process of V(D)J recombination, wherein V and J genes are recombined with nucleotides deleted from their ends and quasi-random non-germline encoded nucleotides inserted at the joining sites. In *β* chains, a D gene is additionally inserted between V and J, but hard to align due to its short length and the CDR3 deletions. We hypothesized that the latent space of TCR-VALID could be able to disentangle V and J gene usage and the mean physicochemical properties of the quasi-random insert region between the V and J genes. Given that the training objectives of TCR-VALID do not explicitly aim to disentangle these predetermined generative factors they likely encode factors beyond those we study, particularly properties relating to the non-mean properties of the insert region.

A well disentangled representation would encode distinct generative factors in non-overlapping subsets of dimensions. To quantitatively benchmark TCR-VALID’s performance and tune its hyper-parameters, we employed a disentangling score [28] that has been identified as robust and broadly applicable [32]. Importantly, this scoring scheme accounts for generative factors that are unknown, but still encoded within the latent space. Briefly this scoring scheme measures the importance of individual latent dimensions for predicting exclusively a single generative factor combined with a weight term accounting for total contribution of the latent term to generative factor prediction (details in methods).

We trained random forest (RF) models [33] on the previously trained 𝒞*β*-VAE latent representations from the sub-sampled TRB training set with a range of capacities 𝒞 to predict the three key TCR generative factors for the associated TCRs (Fig.3a, methods). The feature importances can then be used to both score the disentanglement (Fig.3d) and be visualized via Hinton diagrams showing which dimensions encode differing generative factors and how strongly (Fig.3c). TCRs projected into the latent dimensions for TCR-VALID (𝒞=2) that encode primarily V gene or J gene usage show clear structural stratification by V gene in these dimensions without any further dimension reduction (Fig.3c, top), and those same features are “mixed” in latent dimensions not identified as encoding V/J genes and encode still unknown TCR factors (Fig.3c, bottom). The TCR-VALID model that displays both the highest disentangling score and lowest reconstruction loss was the same model that presented the smoothest learned landscape with capacity of 2 nats per dimension. One potential flaw in the disentangling score, is that it aggregates all generative factors into a single score [32]. We found via hyper-parameter tuning that capacity 2 gives the best balance between disentangling score and sequence reconstruction accuracy as captured by the reconstruction loss (Fig.3d), whilst PCA and autoencoder models sacrifice reconstruction accuracy or latent landscape disentangling.

**FIG. 3.**
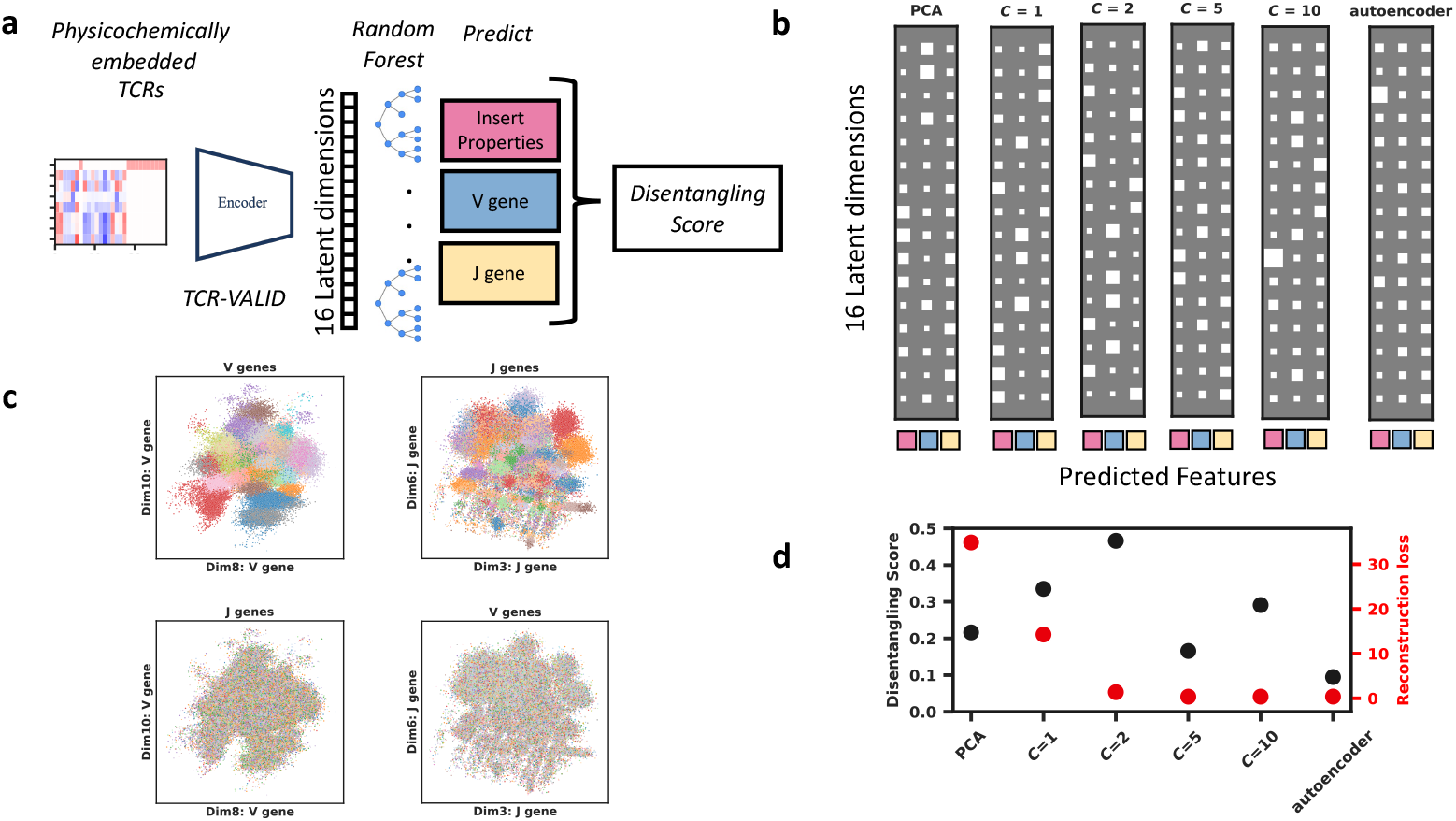
TCR-VALID learns a disentangled interpretable latent landscape. (a) Quantitatively evaluating the disentangling of TCR-VALID by training random forests to predict TCR-intrinsic factors in the learned latent dimension and computing a Disentangling Score [28] on their feature importances. (b) Hinton diagrams illustrating the relative importance of latent dimensions from trained networks at predicting TCR intrinsic features. Square size indicates relative importance, rows indicate latent dimensions and columns indicate respectively mean insert physicochemical properties, V gene and J gene. (c) Plotting the learned latent landscapes with the feature they best encode (top) show a smooth yet disentangled landscape for V (Left) and J (right) genes. If we plot the none encoded feature in the inappropriate dimensions the organization is lost (bottom) d) Disentangling score versus reconstruction loss for TCR-VALID hyperparameter range versus PCA and autoencoder approaches.

### Unsupervised landscape provides interpretable and generalizable antigen specific TCR clustering

TCR sequences that are projected closely into the latent landscape of TCR-VALID have closely related physicochemical features which might underlie their binding mechanisms to the same antigen peptides [34]. We sought to benchmark the TCR-antigen clustering capabilities of TCR-VALID against current state of the art approaches. We fixed 𝒞 = 2, determined to give us the optimal smoothness and disentangling properties and trained a TCR-VALID model on the full training dataset of approximately 100 million unique TRA and TRB chains (methods: data) in order to evaluate its TCR-Antigen clustering capabilities.

We used a comprehensive labeled TCR-antigen dataset (methods) of paired-chain TCRs that bind a wide range of antigens to evaluate our model’s generalizability. We devised a model agnostic method of TCR-antigen clustering scoring to benchmark existing tools fairly on the distance graphs they build. Briefly we use a density based soft clustering approach [35] that has a radius hyperparameter *ϵ* to tune to sensitivity of the clustering. For the iSMART method [36] we tune the related internal distance parameter ‘threshold’. The benchmarking uses two parameters: clustering precision that measures the percentage of clustered TCRs in ‘pure’ clusters of at least 3 unique TCRs (methods) and a clustering Critical Success Index (CSI) that measures the percentage of TCRs of all TCRs that are in ‘pure’ clusters (See methods). We chose a minimal cluster sizes of 3 TCRs as this is a common use-case for TCR clustering, where experimental samples often only capture a small number of of functionally related unique TCRs across different samples. We equally benchmarked clustering with minimal cluster size 2 (Supplementary Fig.2) as is default for iSMART.

We find that TCR-VALID performs as well as sequence-based approaches designed in part by human guided feature selection such as tcr-dist [37] and iSMART [36] in both clustering precision and CSI (Fig.4a). Further, we benchmarked TCR-VALID against recent general protein transformer-based models [20] and TCR specific transformer models [11] (Fig.4b) that learn high dimensional embeddings of TCR sequences (methods) and found our approach significantly outperforms these approaches (Full benchmarking with matched features and chains in Supplementary Fig.1). TCR-VALID’s disentangled representation allows us to probe which dimensions, and associated quantities, independent clusters of TCRs that bind the same antigen differ. We find that the two largest clusters (large cluster n=363, small cluster n=40 unique sequence) we identify from with TCR-VALID bind A*02:01 GILGFVFTL (influenza) peptide with differing motifs as were previously identified in this dataset with a supervised approach [3]. The representative TCR sequence for the binding cluster is generated from averaged TCR-VALID representation which is first decoded back into physicochemical property via the trained decoder and then converted into probability of amino acid identity (Fig.4c) (Details in Methods). This highlights the potential for *de novo* TCR design within region of interest with the TCR-VALID landscape.

**FIG. 4.**
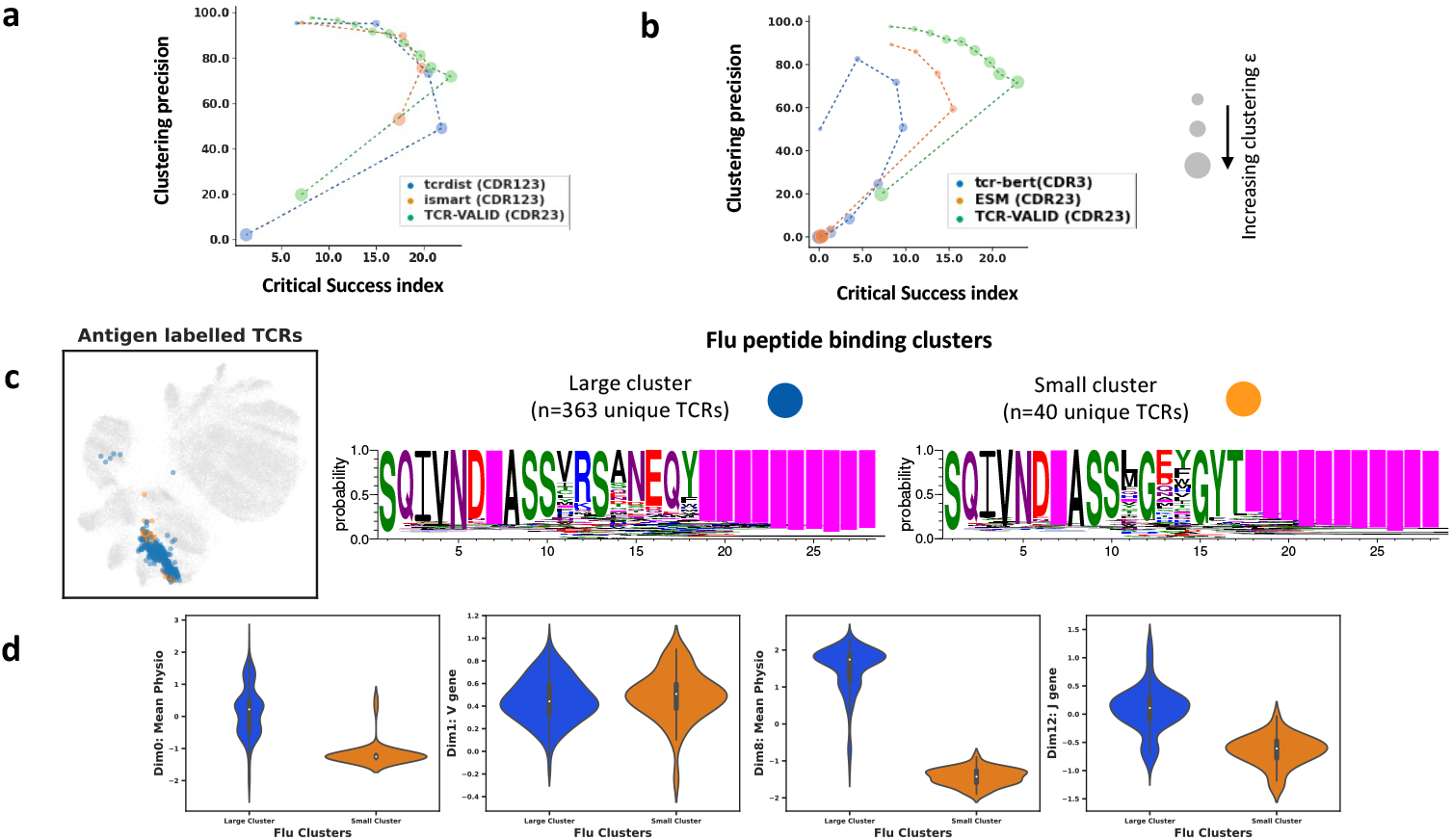
TCR-VALID clusters TCR-antigen pairs and reveals interpretable insight into binding modes. (a) Clustering precision versus Clustering-Critical Success Index (details in methods) for TCR-VALID versus TCR sequence based distance graphs. Increasing circle radius indicates increasing range of DBSCAN *ϵ* radius (b) TCR-VALID vs Transformer based networks. (c) TCRs from validated TCR-antigen pairs embedded into the learned latent landscape (left) and TCRs from two largest flu clusters embedded into learned latent landscape. (c, right) TCR PWMs generated from the mean TCR-VALID representations of the two largest flu clusters, illustrating conserved CDR3 motif and gene usage. (d) Latent dimensions of TCR-VALID model for two flu clusters above illustrating separation of clusters along Mean Insert Physicochemical properties and J gene usage but overlapping for V gene usage.

We confirm that the two clusters share V gene usage (TRBV19) but differ in J gene usage. As expected, the insert region is variable both within a cluster and between clusters. As validation of the disentangling and interpretability of our learned latent landscape, we showed that in dimensions that encode mean insert physicochemical properties (0,8) and J gene usage (12) the two clusters clearly segregate. Conversely, given that the clusters largely share V gene usage, along dimension 1 that encodes V gene we find the two clusters overlap (Fig.4d). These results highlight the benefits of the interpretable learned landscape when relating TCR-antigen binding clusters in an unsupervised fashion to both recover known TCR features such as V,J gene usage and learn biophysical properties such as mean physicochemical insert value which underlie different binding mechanisms.

### Latent representations provide universal feature extractor for TCR classification with uncertainty-estimation

TCR clustering is particularly useful in extracting putative groups of functionally similar TCRs in an antigen agnostic way, however there are some antigens for which many cognate TCRs are known. In those cases, we can build meaningful classification models by learning antigen specific non-linear mappings from the TCR sequences, rather than relying purely on proximity in a physicochemical latent space [3]. This can be particularly useful for common viral antigens, which can be present in large numbers in T cell samples due to clonal expansion. It can be useful to remove these from downstream analysis or even to understand the role of bystander TCRs for instance in immune-oncology settings [38–40].

One problem with classification models is how the models behave on data that is out-of-distribution (OOD). This is problematic in biological data settings [41, 42], in particular for TCRs because of the many possible classes (e.g antigen labels) and only observing a small portion of the input space that belong to specific classes [43]. The simplest route to OOD detection is to consider that samples with a more uniform prediction over the classes is more likely to be OOD, although in practice this is not very performant on modern neural nets with diverse biological data [41].

To improve our OOD detection we utilize not only our labeled dataset for classification, but additionally a portion of the unlabeled data. We use the method of Lee et al. [44] to add an auxiliary loss to force the classification probabilities to the antigen targets to be more uniform when samples come from the unlabeled set, and the usual cross-entropy for the labeled data (methods). These loss terms are balanced by a parameter *α*. Tuning *α* from zero, where there is no OOD detection improvement included, to larger values of *α* where the output distribution is progressively forced to be uniform for unlabeled data. We find that *α* = 1 minimally effects classification AUROC compared to *α* = 0 while improving our ability to classify new TCR data as in-distribution or OOD from AUROC from 0.737 to 0.838 (Table.I). This allows users to make predictions of whether newly observed TCRs bind any in a set of antigens for which some amount of TCR-antigen data was available. Simultaneously users obtain a predictive score of whether the new TCR in question is in-distribution for the model, aiding users identify potential false positives.

**TABLE I.**
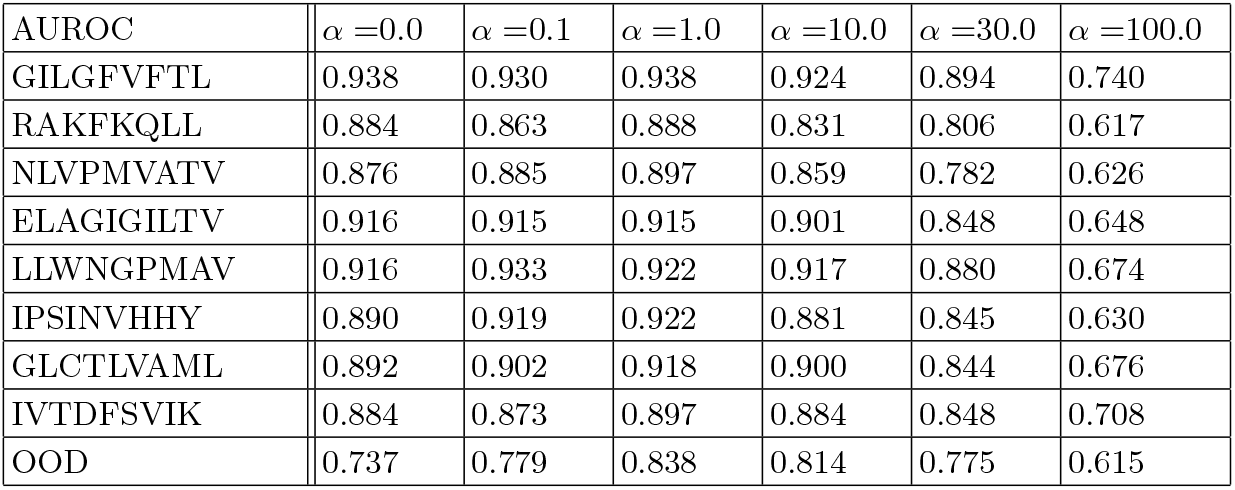
Out of distribution TCR-antigen benchmarking. Per peptide classification AUROC at various out of distribution *α* weighting using a small multilayer perceptron network directly on TCR-VALID representations. The out of distribution AUROC in the final row shows that by tuning *α* one can improve OOD detection while retaining peptide classification AUROC.

## DISCUSSION

In this study we present TCR-VALID, a capacity controlled *β* VAE trained using approximately 100 million unique TCR sequences to learn smooth, interpretable, reduced dimensional representations of TCRs. TCR-VALID is fast, lightweight, and provides state of the art clustering of TCRs by antigen binding properties in its learned latent space without need for retraining.

Other auto-encoding frameworks have been applied to TCR data [5, 13, 24, 45], they have previously either been limited to antigen-labeled TCR data thereby limiting the scope of TCR sequence diversity captured in the representations, have not investigated the disentangled nature in a quantitative manner, or have not investigated the clustering capabilities within the latent space. We applied robust validated metrics from the disentangling representation learning literature [32] and quantitatively assessed our learned latent space smoothness, to optimally choose the information bottleneck of TCR-VALID. To the best of our knowledge this is the most comprehensive investigation of TCR sequence latent space using the quantitative tools of DRL. As a result, TCR-VALID allows us to generate realistic TCR sequences along interpretable dimensions whilst clustering TCR-antigen pairs in an unsupervised fashion as well as current state of the art approaches.

One future use of TCR-VALID’s disentangled TCR physicochemical landscape is expected to be for circumventing some of the current limitations in TCR repertoire profiling. Namely, an inability to relate unique yet similar TCRs seen across individuals. This is often the case in disease settings, where due to limited TCR overlap between individuals few TCRs are found shared between individuals despite a shared disease and therefore assumed TCR response. Since we have shown that TCRs binding the same antigens cluster in TCR-VALID’s latent space, TCRs embedded in TCR-VALID’s latent space will cluster together if functionally similar. Clusters of TCRs present in patients with shared disease, or under the same treatment, may provide a tool to reveal TCRs involved in those conditions, and via the disentangled space a route to understand the conserved nature of those similar TCRs.

Other approaches have shown promise in the DRL literature such as FactorVAE [46] and *β*-TCVAE [47] that further reinforce the disentangling objective of *β*-VAE by adding further terms to the training loss based on latent dimension correlations and an additional discriminator respectively. These tools may provide an increase to the disentangling abilities of TCR representations. In this work we have shown that the key generative factors for TCR sequences, can be well disentangled by a capacity controlled-VAE in a small number of dimensions and with good reconstruction accuracy. This low dimensional representation in combination with the smoothness of the latent space may open up possibilities for latent space optimization of TCR sequences, which would be an exciting avenue for future research.

## METHODS

### Data

We collected two sets of TCR data, an unlabeled set of TCRs from repertoire-level data, that is without corresponding antigen-binding information, and a set of antigen-labeled TCRs for which the cognate antigen is known.

We collected repertoire level TCR data from the iReceptor Gateway [48] (https://gateway.ireceptor.org/login) and VDJServer [49] (https://www.vdjserver.org). These data include TRB and TRA chains, that are predominantly unpaired.

We collected paired-chain TCRs with known cognate antigens from two sources; those associated with [3] and VDJdb [50, 51]. For VDJdb we collected all human paired-chain TCRs with a quality ‘score’ of at least 1 (accessed October 2021).

### Data Preparation

The ingested TCR data is in the AIRR schema (https://docs.airr-community.org/en/v1.2.1/datarep/rearrangements.html). We clean the ingested data to train the model on high-quality TCR sequences. The first step in the quality control pipeline is selecting the locus of the TCR, as either TRB or TRA, and chains that are True for ‘productive’ and ‘vj_in_frame’, and False in ‘stop codon’. Several criteria were used to evaluate the quality of the junction sequences of the TCRs. The amino acid junction sequence must have a length greater than or equal to 7, a length less than 24. The junction must start with the amino acid C, end with the amino acid F, and not contain X, *, or U. Very few sample TCRs have the CDR1 and CDR2 sequences labeled. However, a large percentage are labeled with the column named ‘v_call’, which gives the V gene that encodes for the CDR1 and CDR2 sequences. We thus use ‘v_call’ to annotate CDR1 and CDR2. To prevent ambiguous ‘v_call’s affecting the quality of CDR labels we restrict TCR chains to those for which the ‘v_call’ contains a single V gene while allowing for multiple alleles of a single V gene. After filtering ambiguous ‘v_call’s we assume all V alleles are 01 as the CDR2 sequence between alleles of the same V gene are almost always identical. To assign CDR1 and CDR2 to each chain we first retrieve the amino acid sequences for each of the TRV genes from (https://www.imgt.org/genedb/). This database provides sequences for all human TRV genes, including each allele, for each gene we retain only 01 alleles. For each sequence we use ANARCI

1. to apply IMGT numbering, and retrieve the IMGT CDR1 and CDR2 regions for each sequence. These CDR1 and CDR2 definitions for each v gene are then joined into the dataset via the ‘v_call’. TCR chains for which any of: ‘v_sequence_end’, ‘cdr3_end’, ‘j_sequence_start’, ‘cdr3_sequence_end’ are null are removed. We define the insert amino acid sequence as the (in-frame) codons in the cdr3 nucleotide sequence which are encoded by 1 or more non-V/J encoded nucleotide, as determined by AIRR sequence schema fields: ‘v_sequence_end’, ‘j_sequence_start’, ‘cdr3_start’, ‘cdr3_end’.

### Labeled data subsets

When running clustering experiments we tested various different subsets of the data, and used differing features as inputs for clustering. We select whether to use:

- chains: TRB, TRA or paired-chain TCRs
- feature: CDR3, CDR2+3, or CDR1+2+3.

We then subsequently only keep TCRs that bind antigens with at least 3 TCRs in the dataset, keep TCRs with length of less than 28 residues (CDR3,CDR3+2) or 35 (CDR3+2+1), and then removed duplicate TCRs based on the feature and chains that were to be used in an ML model.

### Unsupervised training data sets

We train models in two regimes: in the first case we use a small fraction of the TRB TCR data (hereon called “smallTRB”) to allow us to sweep hyperparameters in a reasonable timeframe and for the final TRA and TRB models we use all of the TRA and TRB data respectively (“large-TRA”, “largeTRB”) for training the models. In both cases we remove duplicated sequences and split the data into train, validation and test sets with sizes 80%,10%,10% respectively. Exact number of unique sequences: largeTRB (94519890), largeTRA (5176669), smallTRB (4253395).

### TCR sequence formats and physicochemical projection

TCR sequences were represented as a combination of CDR regions (CDR3, CDR3+2, or CDR3+2+1). The CDR regions were joined with gap characters, and this was applied to either TRA-alone, TRB-alone to generate a feature for that TCR chain, and for paired-chain format the features for each chain were joined with a gap character.

To embed amino acid sequences of the TCR sequence, represented by the CDR combination being used, into physicochemical space we project using a normalized version of the physicochemical properties for each amino acid as they appear in Table 1 of [53]. Each amino acid is converted into a vector of 8 values, the seven physicochemical properties and a reserved feature indicating if the amino acid is a gap character, and normalize the values using z-score for each feature over the 20 amino acids. We can denote this space as *{****f*** ^*a*^|*a* ∈ amino acids*}*.

After each amino acid in a sequence is converted a sequence is then represented by a 2D array with the length of the original amino acid sequence and width 8. We pad the end of the amino acid sequence with gaps such that all input data have the same array size, maximal size included is indicated in the following section and differs depending on which CDRs are included. This physicochemical encoding for a given sequence can be written as *x* where 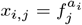 where *a*_*i*_ is the amino acid of the *i*^th^ amino acid in the sequence.

A physicochemical representation of an amino acid sequence can be uniquely projected back to the original sequence, but with any minor alteration to physicochemical property at a given position the amino acid at the position can no longer be uniquely identified and must instead be a probability distribution over the amino acids. This probability assignment is made via a distance metric *δ*(*·,·*) on each physicochemical feature vector and the features of the true amino acids, subsequently normalized to a probability distribution. Namely, for a physicochemical representation 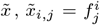, the probability for each amino acid for position *i* is *P* (*a*_*i*_ = *a*|***f*** ^*i*^) = *δ* (***f*** ^*i*^, ***f*** ^*a*^ *)/*Σ_*a*_ *δ* (***f*** ^*i*^, ***f*** ^*a*^). For the distance metric we used *δ* (***f*** ^*i*^, ***f*** ^*a*^) = 1/ (‖ ***f*** ^*i*^ − ***f*** ^*a*^ ‖_1_ + *ε*) with *ε* = 1e-6. This allows any physicochemical feature array to be converted to a PWM over amino acids.

### Capacity-controlled VAE models

#### Architecture

TCR sequences projected into physicochemical arrays, *x*, are fed into an encoder model 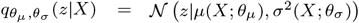 to generate latent space samples *z*, which are subsequently decoded by *p*_*ϕ*_(*X*|*z*).

Since our input sequences are fairly short (≤ 28) we utilize a lightweight 3 layer CNN for our encoder with He normalization and stride of 1 for all 1D convolutions, with 32,64,128 channels and kernel widths 5,3,3. All convolutional layers are followed by Batch Normalization [54] and leaky ReLU activation. Following the the convolution the output is flattened to a vector and two single feed forward layers with output dimension 16 are used to construct the mean and log variance for the sampling of the latent representation to be passed to the decoder. For the decoder a ReLU activated feed forward layer constructs a 28×128 array on which 3 1D deconvolution layers are applied with channels 128,64,32 and kernel widths 3,3,5, and each is followed by Batch Normalization [54] and leaky ReLU. Final reconstructed physicochemical representation is generated by an 8 channel deconvolution layer with kernel width of 1 without activation.

#### Training regime

The loss function is that described by Burgess et al. [26]: ℒ = ℒ_recon_ + *β* | ℒ_KL_ − *n*_*L*_ 𝒞|, where *n*_*L*_ is the number of latent dimensions used and the loss terms used for sample *x* were 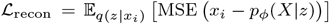 and ℒ_KL_ = *D*_KL_ (𝒩 (*μ*(*x*_*i*_; *θ*_*μ*_), *σ*^2^(*x*_*i*_; *θ*_*σ*_) || 𝒩 (0, 1)). In contrast to Burgess et al. [26] we do not adjust the capacity 𝒞 during training, thereby fixing an average number of nats of information that a dimension of the latent space should aim to encode. We use a value of *β* sufficient to enforce the average capacity is close to the objective, *β* = 1 was sufficient for 𝒞 ≥ 2 whereas *β* = 10 was required for 𝒞 = 1. We note that since MSE loss is used for the reconstruction loss *β* = 1 does not carry the same meaning in terms of the ELBO that it does in the context of a Bernoulli output for binary X, see e.g [55].

We minimize the loss using the Adam optimizer [56] with a learning rate of 1e-3. For the “smallTRB” and “largeTRA” models we use early stopping on the validation data split with a patience of 10 epochs and restore the weights to the epoch at minimal validation loss. For the “largeTRB” model due to the size of data we used checkpointing at 10 epoch intervals and used the checkpointed model at approximate minima of the validation loss.

Training, validation and test datasets were saved in parquet format in either raw sequence format, ingested using Hugging-Face’s [57] “read_parquet” method and converted to physicochemical properties on the fly (for small TRB and largeTRA), or (for largeTRB) TRB sequences were first converted to physicochemical arrays in and saved in parquet format and then ingested using the read parquet method.

### Metrics

#### Clustering Metrics

A ‘good’ cluster is one in which the modal antigen label of TCRs is the label of *>*90% of TCRs in the cluster, following the definition in [58]. If we consider such TCR clones to be ‘clustering true positives’, c-TP, while TCRs in clusters which don’t fit this definition as ‘clustering false positives’, c-FP, and equivalently TCRs that aren’t clustered at all to be ‘clustering false negatives’, c-FN, we can make the following analogies between clustering and classification metrics:

#### c-Precision

Of all clustered TCR clones, the percentage of TCR clones that are clustered into clusters that are ‘good’. Which can be written:

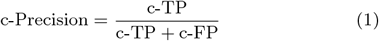

#### c-CSI

The Critical Success Index, of all TCRs how many were placed into ‘good’ clusters:

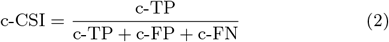

#### Minimal cluster sizes

In order to score the ability of a given TCR distance-metric in combination with a given clustering algorithm to accurately cluster TCRs, we take an approach similar to that of [59] smaller clusters can skew the model performance metrics. We therefore chose to only consider clusters with at least 3 TCRs present. However, we also consider (Supplementary Fig. 2) the case where TCR clusters of only 2 TCRs are allowed and find that physicochemical properties alone perform as well as ismart and tcrdist on TRB chains with CDR123 considered.

#### Continuity Metrics

We project two TCR sequences with identical CDR3s but differing CDR2 into physicochemical space (*x*_1_, *x*_2_) and then into a models latent space (*z*_1_, *z*_2_). We then linearly interpolate between *z*_1_ and *z*_2_ via : 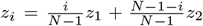 for *i* = 0, 1,, …, *N* − 1. Interpolated latent space representations *z*_*i*_ are then decoded to physicochemical property maps 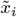.

Inspired by [23] we devise measures of how far interpolated TCRs differ from the true manifold and expected pathway of TCRs between our endpoints. We identify if the CDR2 region of the interpolated TCR remains close to the true manifold of possible TRBV CDR2s and whether the CDR3 region differs significantly from the true CDR3 of both TCR_1_ and TCR_2_.

The distance of an interpolated TCR physicochemical representation to a true TRBV CDR2 can be written:

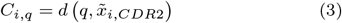

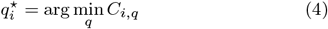

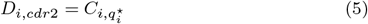

where q is the physicochemical representation of the q^th^ true CDR2 and *d* is a distance metric in the physicochemical space and 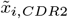 is the CDR2 region of the physicochemical representation 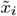. For *d* we use *d*(*x, y*) = Σ_*j*_ ‖***f***^*x*.*j*^ − ***f***^*y*.*j*^ ‖_1_. *D* _*i,cdr*2_ is the distance from an interpolated TCR CDR2 to the nearest true CDR2. We can average this quantity over a trajectory to get a score the the trajectory (smaller is better):

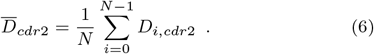

We can also assess how far the CDR3 changed from the true CDR3 at each point on the trajectory:

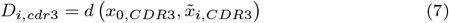

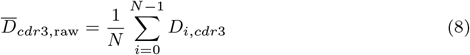

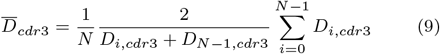

where *x*_0,*CDR*3_ is the physicochemical representation of the CDR3 of the original TCR sequence. 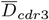 is the mean distance along the trajectory, normalized to the error in reconstruction at the start and end point of the trajectory. This is required to normalize across models with different reconstruction accuracy since we wish to assess changes in the manifold matching, rather than measure reconstruction accuracy which varies as capacity of the model is varied.

To compare models we generate many TCR interpolations and score them via 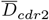 and 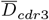. We generate TCR pairs with the same CDR2 but different CDR3, and in particular for each model we generated 310 interpolations - we chose 31 random CDR3s selected from the test set, and for each CDR3 interpolated between 10 random pairs of CDR2s of length 6 amino acids.

#### Disentanglement Metric

We train random forest classifiers (sklearn [60]) with TCR latent space representations as features and V or J gene as labels. In order to create the mean insert physicochemical value we average the physicochemical properties of all the amino acids attributed to the insert region as defined in the data preparation section. This value is then used to train a random forest regressor. In order to evaluate our classifiers we score them using weighted one versus ‘rest’ AUC ROC with stratified 5 fold cross validation. Hyperparameters for the random forest are chosen based on the cross-validation and RFs are retrained with those parameters on the full dataset. For training data for the RFs we use a random selection of 100k TCRs. We then take the random forest feature importances and apply the disentanglement metric as in [28].

### Uncertainty Aware Classification

We used TCR-VALID models to generate representations for the TRB and TRA chains of paired chain TCRs in our labeled dataset restricted to TCRs with binding to antigens for which at least 100 unique TCRs were present in the dataset (8 antigens), and for 100k random unique chains from each of the unlabeled TRA and TRB datasets. We built a feed-forward neural network with the following layers: dropout (25% retention), dense (128, ReLU), dropout (40% retention), dense (128, ReLU), dropout (40 % retention), dense (8, softmax). We constructed input batches of 32 labeled TCRs and 512 randomly selected unlabeled TCRs and trained the neural network with loss function [44]:

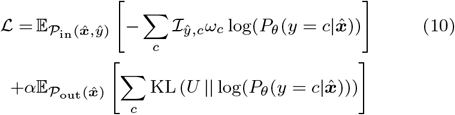

Where we apply weights, *ω*_*c*_, to the cross-entropy loss to combat dataset imbalance, ℐ_*ŷ,c*_ is the indicator function, and *α* is a tunable hyperparameter. *ω*_*c*_ are calculated using sklearn’s ‘compute_class_weight’ function. During training we apply an early stopping criteria with a patience of 5 epochs on the validation set. AUROC for peptide classification are calculated in a one vs rest fashion per peptide on the left out test set. As a measure for detecting OOD we data we calculate *KL* loss on each sample of test data from the labeled set and similarly for the unlabeled set, and subsequently divide KL loss by the maximal *KL* value across all test data. We then use these normalized *KL* scores as effective probabilities for measuring whether data is ID or OOD and use these probabilities to construct an ROC curve and calculate the AUROC for OOD detection. Reported AUROC (Table.I) are calculated as the mean over 5 Monte Carlo cross-validation splits.

### Comparator methods

#### Physicochemical properties

We project the residues of TCRs into their physicochemical properties, constructing a 2D”image” for each TCR as we do for VAE input, and then flatten these 2D images to 1D vectors. These 1D representations are constructed for the subset of labeled TCR data of interest, and clustering is then performed on these representations via Euclidean distance via DBSCAN [35] via scikit-learn’s implementation [60] (similarly for all DBSCAN implementations discussed below).

#### PCA on physicochemical properties

These 1D physicochemical representations are constructed for the subset of labeled TCR data of interest, and for a subset of the unlabeled TCRs. We fit PCA to the unlabeled TCR representations, and project antigen-labeled TCRs into the PCA dimensions. TCR clustering is then performed on PCA representations with DBSCAN. We use inverse PCA transform to project TCR representations in PCA space back into physicochemical space, and subsequently convert from physicochemical space to probability distribution over residues to construct generated TCRs “logos”, via weblogo [61], from any position in the PCA latent space.

#### tcr-dist

We used tcr-dist [37] (https://github.com/phbradley/tcrdist, SHA: 1f5509a, license: MIT) to extract the tcr-dist distances between all pairs of clones in each set of input data. We used this package in two ways. Firstly, using the TCR-TCR distance matrices saved by tcr-dist to employ our own clustering on the resulting graph, in this paper we used DBSCAN and we performed such clustering for TRB-only, TRA-only and paired-chain clustering. tcr-dist also identifies TCR clusters via its own clustering algorithm and we modified the tcr-dist code (specifically `make_tall_trees.py`) to allow us to assess the clustering performance of these clusters on the tcr-dist distance graph. In the original tcr-dist implementation clustered TCRs are saved only as logos in an image (file ending `tall_trees.svg`). Internally tcr-dist finds small TCR clusters, and hierarchically joins them into larger clusters, these larger clusters are those shown in the output image, and we therefore use those cluster definitions here. Additionally, TCR clones are allowed to be in more than one cluster, this does not work with our scoring paradigm, so for TCRs which were assigned to more than one cluster we chose to assign such TCR clones to the cluster with maximal TCR membership.

#### iSMART

We use iSMART [36] (https://github.com/s175573/iSMART, SHA: 1f2cbd2, license: GPL3.0) to cluster TRB-only data, as it was not designed to cluster TRA or paired-chain TCR data. We tune the internal distance parameter, `threshold`, within their clustering method to tune the size of the clusters.

#### TCR-BERT

We use TCR-BERT [11] in two modes, their internal clustering method and DBSCAN clustering applied to a distance graph defined by distance between the TCR-BERT embeddings. TCR-BERT embeddings are calculated from the output of the eighth layer of the transformer model, as used within the original publication due to this layer’s outputs being “optimal” [11]. We use the script `embed_and_cluster.py` to perform tcr-bert clustering, tuning the Leiden resolution to adjust the cluster sizes. It is worth noting that this clustering assigns a cluster to all TCRs, in contrast to DBSCAN which allows TCRs to not belong to any cluster. We also collected the TCR-BERT embeddings from the output of the eighth layer (using ‘mean’ aggregation as per the tcr-bert default) and used Euclidean distance between these embedding representations to build a distance graph on which we could apply DBSCAN clustering.

#### ESM

We used ESM-1b [20] to generate representations of TCR sequences in several ways. We gave the TCR sequences as input with either the CDR3 alone, or the CDR2 and CDR3 joined with a single gap. To generate TCR distances from the TCR representations generated by ESM1b we employ Euclidean distances between the representations, either on the entire representation (flattened to 1D) or the mean-pooled representation. DBSCAN is used as the clustering algorithm on this TCR distance metric.

## DATA AVAILABILITY

Collected repertoire level TCR data from the iReceptor Gateway [48] (https://gateway.ireceptor.org/login) and VDJServer [49] (https://www.vdjserver.org) and is publicly available. A list of the repertoire_id’s of the repertoires used in this study will be included in our github (https://github.com/peterghawkins-regn/tcrvalid). Collected paired-chain TCRs with known cognate antigens from two sources; those associated with [3] and VDJdb [50, 51] (all human paired-chain TCRs with a quality ‘score’ of at least 1 (accessed October 2021)) are also publicly available, the exact dataset will be deposited in our github on peer-review.

## CODE AVAILABILITY

The tcrvalid package, and pre-trained models, are available at https://github.com/peterghawkins-regn/tcrvalid.

## COMPETING INTERESTS

The authors are employees of, and have stock and stock options in, Regeneron Pharmaceuticals Inc. G.A. is an officer of Regeneron Pharmaceuticals Inc.

---

**Supplementary Figure 1.**
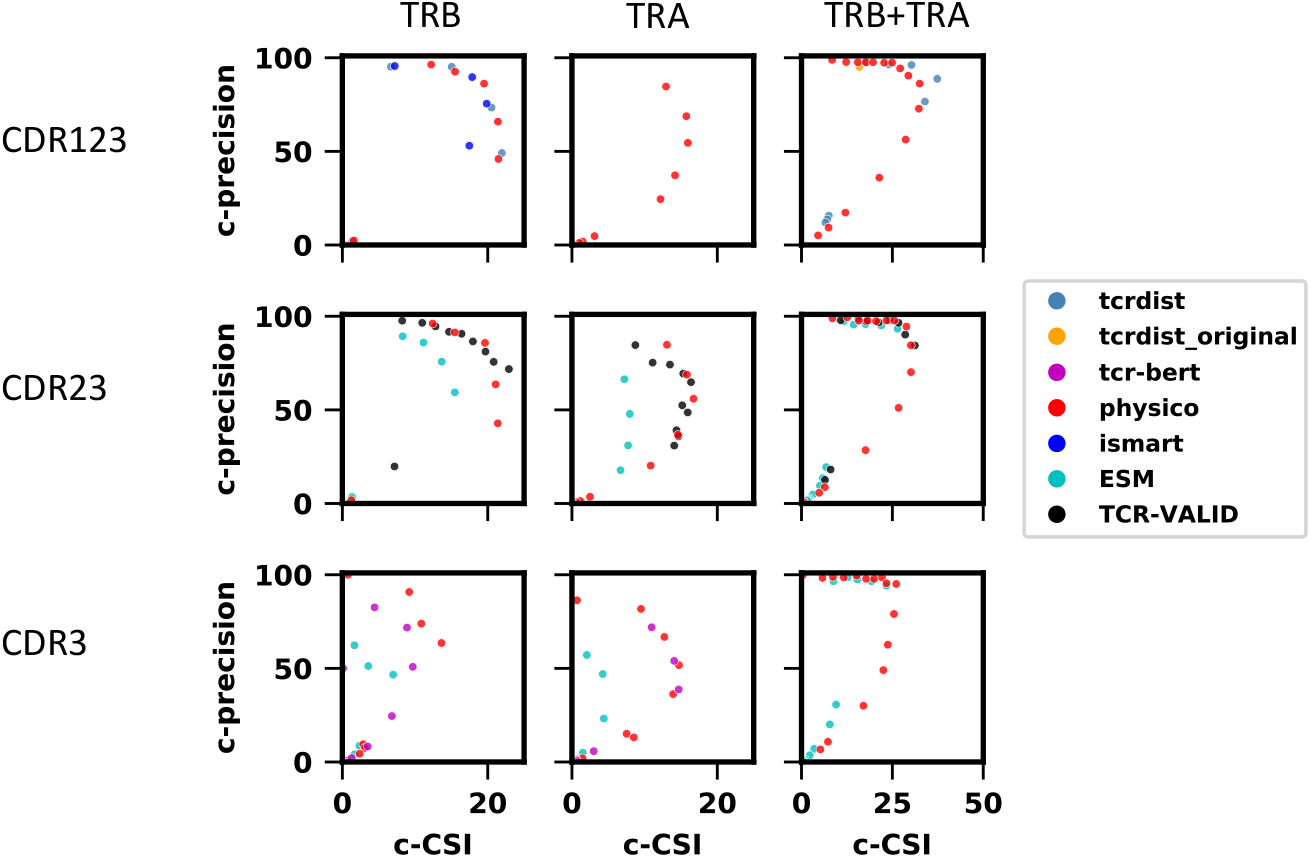
Benchmarking TCR-Antigen clustering algorithms. Clustering precision versus Clustering-Critical Success Index for TCR-VALID versus both sequence based (tcrdist and ismart), transformer based models (tcr-bert and ESM). We additionally benchmarked clustering of TCRs based solely on physicochemically featurized sequences (physico) as those are the base features provided to TCR-VALID. The tcrdist [37] and ismart [36] methods developed their own clustering approach and as such we benchmarked both the published tcrdist (tcrdist-original) method, DBSCAN (tcrdist, min cluster size=3) on the tcrdist distance graphs along with the published ismart method. The columns benchmark single chain and paired chains (TRB, TRA, TRB+TRB respectively) whilst the rows benchmark the CDR regions employed (CDR123, CDR23, CDR3 respectively))

**Supplementary Figure 2.**
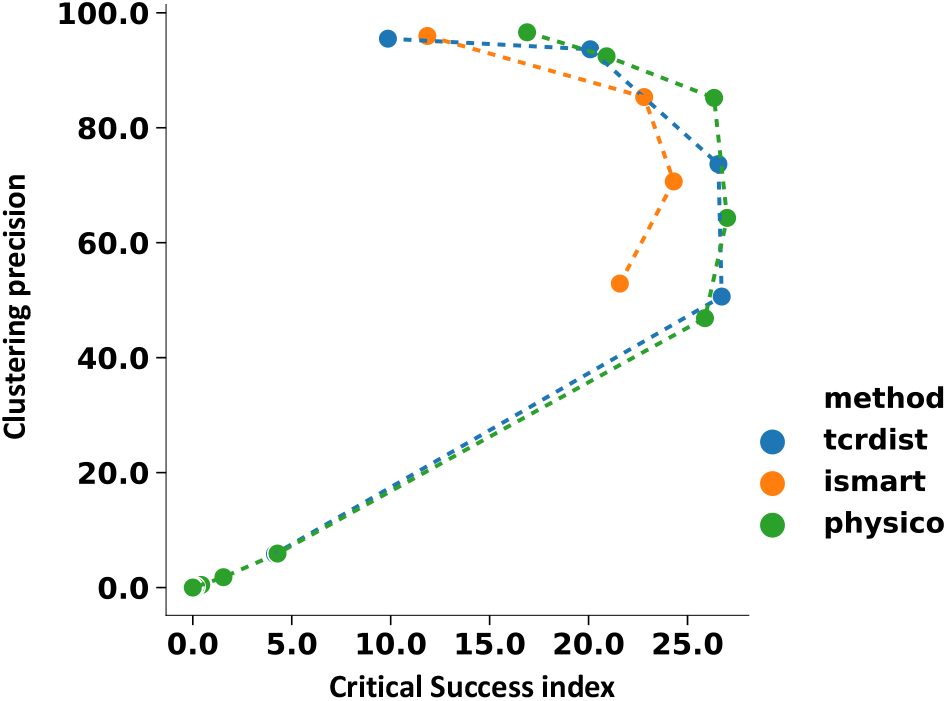
Benchmarking TCR clustering with minimum cluster size=2. Both tcrdist and ismart approaches have a minimum cluster size of 2 but when using DBSCAN we chose minimum cluster size of 3 in order to capture TCR density in TCR-VALID latent space. In order to benchmark we measured Clustering precision versus Clustering-Critical Success Index for minimum cluster size 2 for tcrdist, ismart and physiochemical properties with TRB chains and CDR123. This allows for a fair comparison of the influence of minimum cluster size on clustering with the different methods but the same chains (TRB) and sequence features (CDR123).)

## References

[1] C. A. Janeway, P. Travers, M. Walport, and D. J. Capra, Immunobiology (Taylor & Francis Group UK: Garland Science, 2001).

[2] J. A. Pai and A. T. Satpathy, High-throughput and single-cell t cell receptor sequencing technologies, Nature Methods 18, 881 (2021).

[3] W. Zhang, P. G. Hawkins, J. He, N. T. Gupta, J. Liu, G. Choonoo, S. W. Jeong, C. R. Chen, A. Dhanik, M. Dillon, et al., A framework for highly multiplexed dextramer mapping and prediction of T cell receptor sequences to antigen specificity,Science Advances 7, eabf5835 (2021).

[4] A. A. Minervina, M. V. Pogorelyy, A. M. Kirk, J. C. Crawford, E. K. Allen, C.-H. Chou, R. C. Mettelman, K. J. Allison, C.-Y. Lin, D. C. Brice, et al., Sars-cov-2 antigen exposure history shapes phenotypes and specificity of memory cd8+ t cells, Nature Immunology 23, 781 (2022).

[5] J.-W. Sidhom, H. B. Larman, D. M. Pardoll, and A. S. Baras,DeepTCR is a deep learning framework for revealing sequence concepts within T-cell repertoires., Nature communications 12, 1605 (2020).

[6] G. Lythe, R. E. Callard, R. L. Hoare, and C. Molina-París, How many tcr clonotypes does a body maintain?, Journal of theoretical biology 389, 214 (2016).

[7] T. Mora and A. M. Walczak, Quantifying lymphocyte receptor diversity, in Systems Immunology (CRC Press, 2018) pp. 183–198.

[8] A. K. Sewell, Why must t cells be cross-reactive?, Nature Reviews Immunology 12, 669 (2012).

[9] B. Daniel, K. E. Yost, S. Hsiung, K. Sandor, Y. Xia, Y. Qi, K. J. Hiam-Galvez, M. Black, C. J Raposo, Q. Shi, et al., Divergent clonal differentiation trajectories of t cell exhaustionNature Immunology 23, 1614 (2022).

[10] A. Purcarea, S. Jarosch, J. Barton, S. Grassmann, L. Pachmayr, E. D’Ippolito, M. Hammel, A. Hochholzer, K. I. Wagner, J. H. van den Berg, et al., Signatures of recent activation identify a circulating t cell compartment containing tumor-specific antigen receptors with high avidity, Science Immunology 7, eabm2077 (2022).

[11] K. Wu, K. E. Yost, B. Daniel, J. A. Belk, Y. Xia, T. Egawa, A. Satpathy, H. Y. Chang, and J. Zou, TCR-BERT: learning the grammar of T-cell receptors for flexible antigen-xbinding analyses, bioRxiv 10.1101/2021.11.18.469186 (2021).

[12] I. Springer, H. Besser, N. Tickotsky-Moskovitz, S. Dvorkin, and Y. Louzoun, Prediction of Specific TCR-Peptide Binding From Large Dictionaries of TCR-Peptide Pairs, Frontiers in Immunology 11, 1803 (2020).

[13] K. Davidsen, B. J. Olson, W. S. DeWitt, J. Feng, E. Harkins, P. Bradley, and F. A. Matsen, Deep generative models for T cell receptor protein sequences, eLife 8, e46935 (2019).

[14] P. Nathan, J. C. Hassel, P. Rutkowski, J.-F. Baurain, M. O. Butler, M. Schlaak, R. J. Sullivan, S. Ochsenreither, R. Dummer, J. M. Kirkwood, et al., Overall survival benefit with tebentafusp in metastatic uveal melanoma, New England Journal of Medicine 385, 1196 (2021).

[15] J. W. Park, S. Stanton, S. Saremi, A. Watkins, H. Dwyer, V. Gligorijevic, R. Bonneau, S. Ra, and K. Cho, Propertydag: Multi-objective bayesian optimization of partially ordered, mixed-variable properties for biological sequence design, arXiv preprint arXiv:2210.04096 10.48550/arXiv.2210.04096 (2022).

[16] R. Gómez-Bombarelli, J. N. Wei, D. Duvenaud, J. M. Hernändez-Lobato, B. Sänchez-Lengeling, D. Sheberla, J. Aguilera-Iparraguirre, T. D. Hirzel, R. P. Adams, and A. Aspuru-Guzik, Automatic chemical design using a data-driven continuous representation of molecules, ACS central science 4, 268 (2018).

[17] A. Tripp, E. Daxberger, and J. M. Hernändez-Lobato, Sample-efficient optimization in the latent space of deep generative models via weighted retraining, Advances in Neural Information Processing Systems 33, 11259 (2020).

[18] A. Grosnit, R. Tutunov, A. M. Maraval, R.-R. Griffiths, A. I. Cowen-Rivers, L. Yang, L. Zhu, W. Lyu, Z. Chen, J. Wang, et al., High-dimensional bayesian optimisation with variational autoencoders and deep metric learning, arXiv preprint arXiv:2106.03609 10.48550/arXiv.2106.03609 (2021).

[19] N. Maus, H. T. Jones, J. S. Moore, M. J. Kusner, J. Bradshaw, and J. R. Gardner, Local latent space bayesian optimization over structured inputs, arXiv preprint arXiv:2201.11872 10.48550/arXiv.2201.11872 (2022).

[20] A. Rives, J. Meier, T. Sercu, S. Goyal, Z. Lin, J. Liu, D. Guo, M. Ott, C. L. Zitnick, J. Ma, et al., Biological structure and function emerge from scaling unsupervised learning to 250 million protein sequences, Proceedings of the National Academy of Sciences 118, e2016239118 (2021).

[21] N. S. Detlefsen, S. Hauberg, and W. Boomsma, Learning meaningful representations of protein sequences, Nature communications 13, 1 (2022).

[22] T. Brown, B. Mann, N. Ryder, M. Subbiah, J. D. Kaplan, P. Dhariwal, A. Neelakantan, P. Shyam, G. Sastry, A. Askell, et al., Language models are few-shot learners, Advances in neural information processing systems 33, 1877 (2020).

[23] D. Berthelot, C. Raffel, A. Roy, and I. Goodfellow, Understanding and improving interpolation in autoencoders via an adversarial regularizer, arXiv preprint arXiv:1807.07543 https://doi.org/10.48550/arXiv.1807.07543 (2018).

[24] M. R. Min, T. Li, H. Guo, F. Grazioli, and M. Gerstein, Learning disentangled representations for t cell receptor design (2022).

[25] D. P. Kingma and M. Welling, Auto-Encoding Variational Bayes, arXiv 10.48550/arxiv.1312.6114 2013), 1312.6114.

[26] C. P. Burgess, I. Higgins, A. Pal, L. Matthey, N. Watters, G. Desjardins, and A. Lerchner, Understanding disentangling in β-VAE, arXiv 10.48550/arxiv.1804.03599 (2018), 1804.03599.

[27] W. K. Wong, J. Leem, and C. M. Deane, Comparative Analysis of the CDR Loops of Antigen Receptors, Frontiers in Immunology 10, 2454 (2019).

[28] C. Eastwood and C. K. I. Williams, A framework for the quantitative evaluation of disentangled representations, in International Conference on Learning Representations (2018).

[29] H. Shao, S. Yao, D. Sun, A. Zhang, S. Liu, D. Liu, J. Wang, and T. Abdelzaher, Controlvae: Controllable variational autoencoder, in International Conference on Machine Learning (PMLR, 2020) pp.8655–8664.

[30] H. Fu, C. Li, X. Liu, J. Gao, A. Celikyilmaz, and L. Carin, Cyclical annealing schedule: A simple approach to mitigating kl vanishing, arXiv preprint arXiv:1903.10145 10.48550/arXiv.1903.10145 (2019).

[31] S. Chakraborty, R. Tomsett, R. Raghavendra, D. Harborne, M. Alzantot, F. Cerutti, M. Srivastava, A. Preece, S. Julier, R. M. Rao, et al., Interpretability of deep learning models: A survey of results, in 2017 IEEE smartworld (smartworld/SCALCOM/UIC/ATC/CBDcom/IOP/SCI) (IEEE, 2017) pp.1–6.

[32] M.-A. Carbonneau, J. Zaïdi, J. Boilard, and G. Gagnon, Measuring disentanglement: A review of metrics, IEEE Transactions on Neural Networks and Learning Systems, 1 (2022).

[33] L. Breiman, Random Forests, Machine Learning 45, 5 (2001).

[34] A. Volkamer, D. Kuhn, F. Rippmann, and M. Rarey, Predicting enzymatic function from global binding site descriptors, Proteins: Structure, Function, and Bioinformatics 81, 10.1002/prot.24205 (2013).

[35] M. Ester, H.-P. Kriegel, J. Sander, and X. Xu, A Density-Based Algorithm for Discovering Clusters in Large Spatial Databases with Noise, in Proceedings of the Second International Conference on Knowledge Discovery and Data Mining, KDD’96 (AAAI Press, 1996)pp. 226–231.

[36] H. Zhang, L. Liu, J. Zhang, J. Chen, J. Ye, S. Shukla, J. Qiao, X. Zhan, H. Chen, C. J. Wu, et al., Investigation of Antigen-Specific T-Cell Receptor Clusters in Human Cancers, Clinical Cancer Research 26, 1359 (2020).

[37] P. Dash, A. J. Fiore-Gartland, T. Hertz, G. C. Wang, S. Sharma, A. Souquette, J. C. Crawford, E. B. Clemens, T. H. Nguyen, K. Kedzierska, et al., Quantifiable predictive features define epitope-specific T cell receptor repertoires, Nature 547, 89 (2017).

[38] Y. Simoni, E. Becht, M. Fehlings, C. Y. Loh, S.-L. Koo, K. W. W. Teng, J. P. S. Yeong, R. Nahar, T. Zhang, H. Kared, et al., Bystander cd8+ t cells are abundant and phenotypically distinct in human tumour infiltrates, Nature 557, 575 (2018).

[39] W. Scheper, S. Kelderman, L. F. Fanchi, C. Linnemann, G. Bendle, M. A. de Rooij, C. Hirt, R. Mezzadra, M. Slagter, K. Dijkstra, et al., Low and variable tumor reactivity of the intratumoral tcr repertoire in human cancers, Nature medicine 25, 89 (2019).

[40] S.-H. Chiou, D. Tseng, A. Reuben, V. Mallajosyula, I. S. Molina, S. Conley, J. Wilhelmy, A. M. McSween, X. Yang, D. Nishimiya, et al., Global analysis of shared t cell specificities in human non-small cell lung cancer enables hla inference and antigen discovery, Immunity 54, 586 (2021).

[41] J. Ren, P. J. Liu, E. Fertig, J. Snoek, R. Poplin, M. Depristo, J. Dillon, and B. Lakshminarayanan, Likelihood ratios for out-of-distribution detection, Advances in neural information processing systems 32, 10.48550/arXiv.1906.02845 (2019).

[42] S. Fort, J. Ren, and B. Lakshminarayanan, Exploring the limits of out-of-distribution detection, Advances in Neural Information Processing Systems 34, 7068 (2021).

[43] A. Weber, J. Born, and M. Rodriguez Martínez, Titan: T-cell receptor specificity prediction with bimodal attention networks, Bioinformatics 37, i237 (2021).

[44] K. Lee, H. Lee, K. Lee, and J. Shin, Training confidence-calibrated classifiers for detecting out-of-distribution samples, arXiv preprint arXiv:1711.0932 10.48550/arXiv.1711.09325 (2017).

[45] T. Lu, Z. Zhang, J. Zhu, Y. Wang, P. Jiang, X. Xiao, C. Bernatchez, J. V. Heymach, D. L. Gibbons, J. Wang, L. Xu, A. Reuben, and T. Wang, Deep learning-based prediction of the t cell receptor–antigen binding specificity, Nature Machine Intelligence 3, 864 (2021).

[46] H. Kim and A. Mnih, Disentangling by factorising (2018).

[47] R. T. Q. Chen, X. Li, R. Grosse, and D. Duvenaud, Isolating sources of disentanglement in variational autoencoders (2018).

[48] B. D. Corrie, N. Marthandan, B. Zimonja, J. Jaglale, Y. Zhou, E. Barr, N. Knoetze, F. M. Breden, S. Christley, J. K. Scott, et al., iReceptor: a platform for querying and analyzing antibody/B-cell and T-cell receptor repertoire data across federated repositories, Immunological reviews 284, 24 (2018).

[49] S. Christley, W. Scarborough, E. Salinas, W. H. Rounds, I. T. Toby, J. M. Fonner, M. K. Levin, M. Kim, S. A. Mock, C. Jordan, et al., VDJServer: a cloud-based analysis portal and data commons for immune repertoire sequences and rearrangements, Frontiers in immunology 9, 976 (2018).

[50] M. Shugay, D. V. Bagaev, I. V. Zvyagin, R. M. Vroomans, J. C. Crawford, G. Dolton, E. A. Komech, A. L. Sycheva, A. E. Koneva, E. S. Egorov, et al., VDJdb: a curated database of T-cell receptor sequences with known antigen specificity, Nucleic acids research 46, D419 (2018).

[51] D. V. Bagaev, R. M. Vroomans, J. Samir, U. Stervbo, C. Rius, G. Dolton, A. Greenshields-Watson, M. Attaf, E. S. Egorov, I. V. Zvyagin, et al., VDJdb in 2019: database extension, new analysis infrastructure and a T-cell receptor motif compendium,Nucleic Acids Research 48, D1057 (2020).

[52] J. Dunbar and C. M. Deane, ANARCI: antigen receptor numbering and receptor classification,Bioinformatics 32, 298 (2016).

[53] J. Meiler, M. Müller, A. Zeidler, and F. Schmäschke, Generation and evaluation of dimension-reduced amino acid parameter representations by artificial neural networks, Molecular modeling annual 7, 360 (2001).

[54] S. Ioffe and C. Szegedy, Batch normalization: Accelerating deep network training by reducing internal covariate shift, in International conference on machine learning PMLR, 2015) pp. 448–456.

[55] C. Doersch, Tutorial on variational autoencoders, arXiv preprint arXiv:1606.0590810.48550/arXiv.1606.05908 (2016).

[56] D. P. Kingma and J. Ba, Adam: A method for stochastic optimization, arXiv preprint arXiv:1412.6980 10.48550/arXiv.1412.6980 (2014).

[57] T. Wolf, L. Debut, V. Sanh, J. Chaumond, C. Delangue, A. Moi, P. Cistac, T. Rault, R. Louf, M. Funtowicz, et al., Huggingface’s transformers: State-of-the-art natural language processing,arXiv preprint arXiv:1910.03771 10.48550/arXiv.1910.03771 (2019).

[58] H. Huang, C. Wang, F. Rubelt, T. J. Scriba, and M. M. Davis, Analyzing the Mycobacterium tuberculosis immune response by T-cell receptor clustering with GLIPH2 and genome-wide antigen screening, Nature biotechnology 38, 1194 (2020).

[59] S. Valkiers, M. V. Houcke, K. Laukens, and P. Meysman, ClusTCR: a python interface for rapid clustering of large sets of CDR3 sequences with unknown antigen specificity, Bioinformatics 37, 4865 (2021).

[60] F. Pedregosa, G. Varoquaux, A. Gramfort, V. Michel, B. Thirion, O. Grisel, M. Blondel, P. Prettenhofer, R. Weiss, V. Dubourg, J. Vanderplas, A. Passos, D. Cournapeau, M. Brucher, M. Perrot, and E. Duchesnay, Scikit-learn: Machine learning in Python, Journal of Machine Learning Research 12, 2825 (2011).

[61] G. E. Crooks, G. Hon, J.-M. Chandonia, and S. E. Brenner, Weblogo: a sequence logo generator, Genome research 14, 1188 (2004).

